# Amyloid fibril structures link CHCHD10 and CHCHD2 to neurodegeneration

**DOI:** 10.1101/2024.07.18.604174

**Authors:** Guohua Lv, Nicole M. Sayles, Yun Huang, Chiara D. Mancinelli, Kevin McAvoy, Neil A. Shneider, Giovanni Manfredi, Hibiki Kawamata, David Eliezer

## Abstract

CHCHD10 is mutated in rare cases of FTD and ALS and aggregates in mouse models of disease. Here we show that the disordered N-terminal domain of CHCHD10 forms amyloid fibrils and report their cryoEM structure. Disease-associated mutations cannot be accommodated by the WT fibril structure, while sequence differences between CHCHD10 and CHCHD2 are tolerated, explaining the co-aggregation of the two proteins and linking CHCHD10 and CHCHD2 amyloid fibrils to neurodegeneration.

## Main Text

Coiled-coiled-helix-coiled-coiled-helix domain containing 10 (CHCHD10, D10) is the first mitochondrial protein to be associated with familial frontotemporal dementia (FTD) and amyotrophic lateral sclerosis (ALS)^1,2^. Numerous pathogenic D10 variants have been reported^3^, including the first mutation to be identified, p.S59L^1^. D10 forms inclusions in disease brains^4^ and mutant p.S55L (equivalent of human p.S59L) D10 knock in (KI) mice (D10^S55L^) develop toxic protein aggregates in multiple tissues, resulting in neurological defects, myopathy, and cardiomyopathy^5-7^, but the nature and structure of these aggregates remain unknown.

The normal function of D10 is unknown, but the protein associates with the mitochondrial inner membrane^8^, its paralog protein CHCHD2 (D2), and other mitochondrial proteins^*8-10*^. Ablation of D10^5,11^ or D2^12^ in mice does not impair mitochondrial functions and has no pathological consequences, while knockout (KO) of both D10 and D2 causes mitochondrial alterations^11^, but very mild phenotypes, suggesting that the proteins can functionally complement each other and are not essential for life. Given the autosomal dominant nature of disease-associated D10 mutations, these observations support a toxic gain of function mechanism.

The N-terminal region of D10 (residues 1-99), lacking the folded CHCH domain, resembles low complexity domains that form the core of amyloid aggregates of other FTD/ALS-associated proteins such as TDP-43, FUS and hnRNPA1^13-17^, and most D10 disease-associated mutations, including p.S59L and p.R15L, fall within this region^18^. A purified recombinant fragment corresponding to this region (D10-NT) is disordered in solution (**Figs. 1A and S1**) and forms amyloid fibrils in vitro when agitated in solution, as judged by Thioflavin T fluorescence (**Fig. 1B**) and TEM (**Fig. 1C**). D10 deposits in skin fibroblasts from patients with the p.R15L disease-associated mutation stain for Thioflavin S, indicating that they contain amyloid aggregates (**Figs. 1D**). The more extensive Thioflavin S staining around the D10 deposits may reflect co-deposition with other amyloids^19^. D10-immunogold staining of sarkosyl insoluble material from D10^S55L^ mice reveals gold-labeled fibrillar species consistent with amyloid fibrils (**Figs. 1E and S2**), which are absent from wildtype (D10^WT^) or D10^KO^ mouse material.

**Figure 1:**
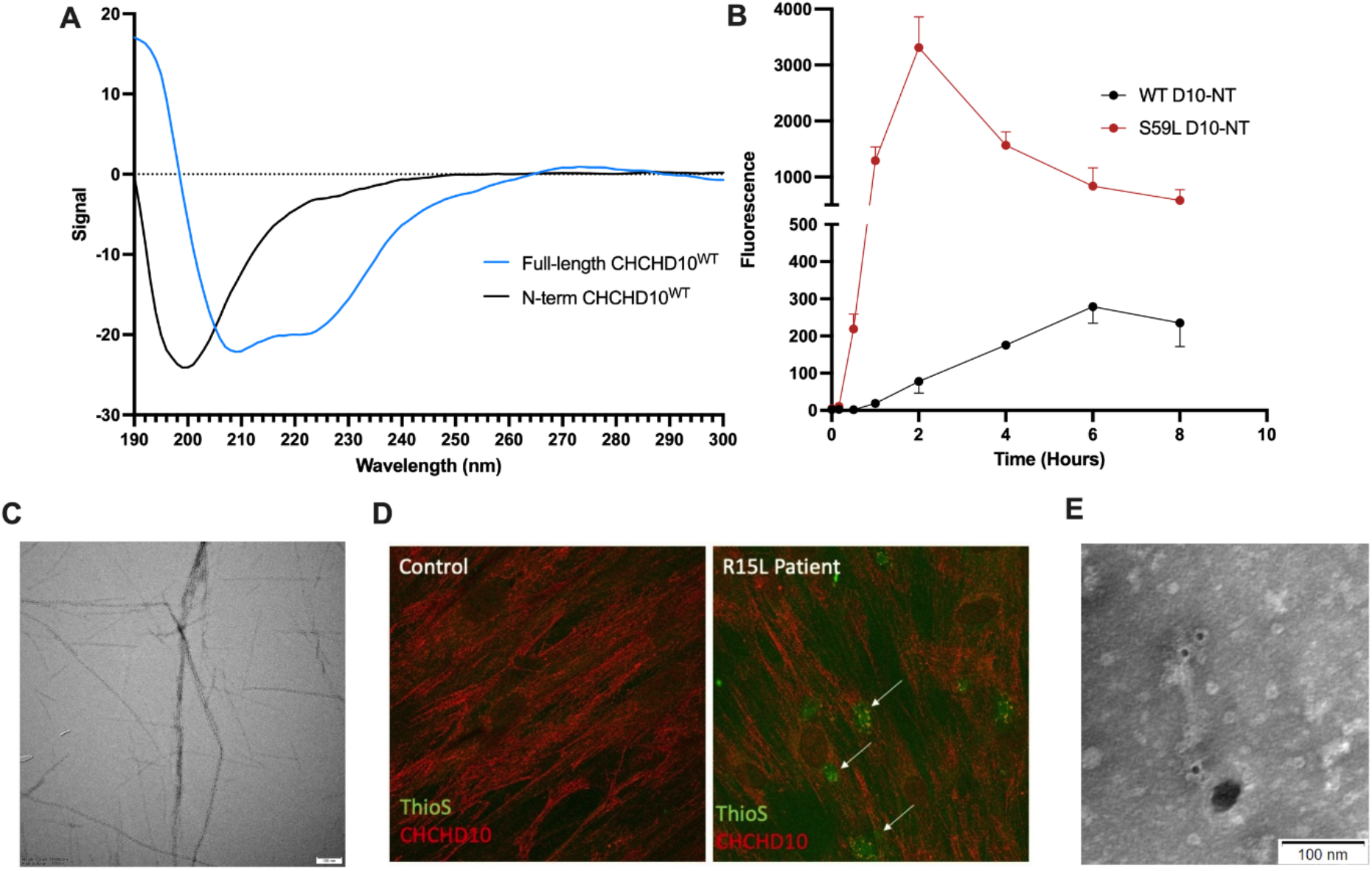
**A**) Circular dichroism spectra of full-length D10 (blue) and D10-NT (black). **B**) Thioflavin T monitored aggregation of purified recombinant D10-NT. **C**) TEM micrograph of aggregated D10-NT. **D**) Skin fibroblasts from p.R15L D10 patients immunostained for D10 (red) and co-stained with Thioflavin S (green) showing colocalization (arrows) **E**) TEM micrograph of D10-immunogold labeled sarkosyl insoluble fraction of heart tissue from a D10^S55L^ mouse.

We determined the structure of D10-NT fibrils using cryo-EM (**Figs. 2A,B and S3**) to a resolution of 2.3 Å (**Fig. 2C**,**S3C**). The ordered core of the fibrils is formed by residues P42-G75 and contains two identical protofilaments (PFs) related by pseudo-2_1_ helical screw symmetry. Each PF contains three beta-strands (**Fig. 2D**), extending from residues M45 to A53, V55-V57 and V64-A72. Strand-2 forms part of a beta-helix like turn structure, commonly observed in amyloid fibrils, connecting the first and third strands. The interface between the PFs is formed primarily between the tip of each beta-helix and the N-terminal half of strand-1.

**Figure 2:**
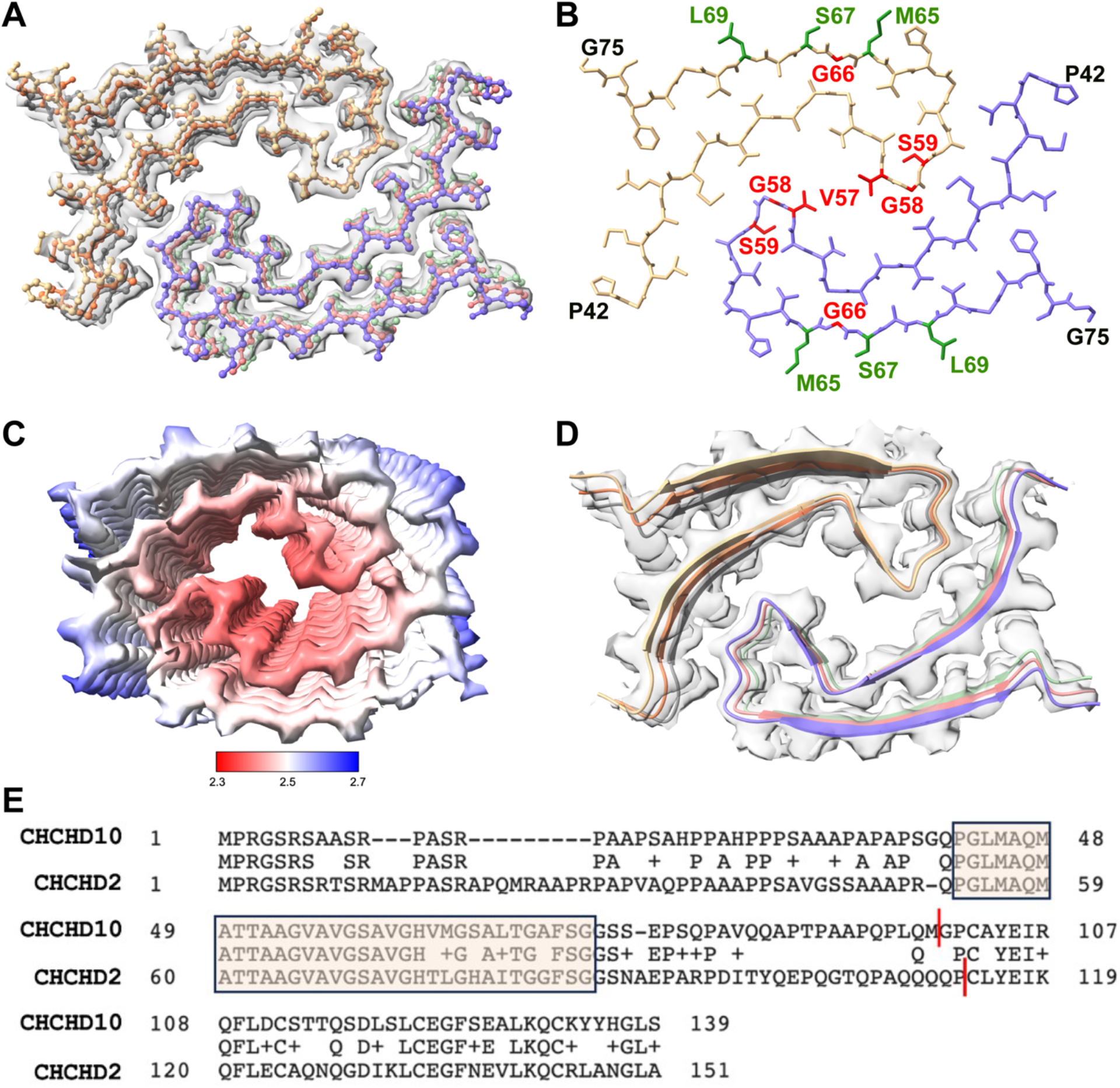
**A**) Cryo-EM density map and atomic model of D10-NT fibrils. Three layers of the fibril structure are shown. **B**) Stick representation of a single layer of the D10-NT ordered fibril core (P42-G75). Disease linked mutations (red) S59L and G66V are situated in the interior of the protofilament fold, and V57E and G58R in the protofilament interface. In contrast, amino acids that differ between D10 and D2 (green) are exposed on the surface of D10-NT fibrils. **C**) Cryo-EM density map colored according to local resolution. **D**) Cartoon representation of secondary structure in the D10-NT fibril core. **E**) Sequence alignment of human D10 (top) and D2 (bottom) showing identical amino acids (middle), conservative substitutions (+) and non-conservative substitutions (blanks). The fibril core is boxed and shaded. Red lines indicate the C-termini of the N-terminal constructs used in our studies.

A number of disease-associated D10 mutations fall within the fibril core region, including p.V57E, p.G58R, p.S59L and p.G66V. Interestingly, S59 and G66 both have small sidechains that are packed in the PF interior (**Fig. 2B**). Clearly, disease-associated substitutions of either of these residues for large sidechains would be inconsistent with the WT fibril structure and would require considerable rearrangements. Residues V57 and G68 are found at the PF interface (**Fig. 2B**), and bulky charged sidechains at these positions would be inconsistent with the packing arrangement of the WT PFs. Thus, disease-associated D10 mutations will likely result in different fibril forms that may underlie the different phenotypes associated with each mutation and constitute structural disease strains, as observed for other amyloids^20,21^. To test this, we solved the structure of S59L D10-NT fibrils (**Fig. S4**). Notably, the S59L mutant aggregates much more rapidly than the WT protein (**Fig. 1B**), providing a potential explanation for its pathogenicity. As expected, residue L59 in this structure flips away from the PF interior, resulting in different PF structures and packing arrangement (**Fig. S4E)**. S59L fibrils are no longer symmetric, with each PF adopting a different fold. PF-1 adopts a fold similar to the WT PFs, except that the beta-helix connecting strands 1 and 3 is disrupted by the flipping of residue 59. PF-2, however, adopts a completely different fold, likely driven by the fact that the WT arrangement could not be maintained with bulky L59 at the PF interface. These results confirm that disease mutations alter D10 fibril structure.

D10 has been reported to co-aggregate with D2, which is also associated with neurodegeneration^5,7^. Alignment of the human D10 and D2 sequences reveals that that the D10 fibril core represents the most highly conserved region between the two proteins, with only 5 sequence alterations (**Fig. 2E**). Three of these alterations (M65L, S67H and L69I) occur at positions facing the outside of the fibril structure (**Fig. 2B**), while the remaining two are isosteric (V64T) or reduce the sidechain size (A72G) and would therefore be expected to be tolerated by the D10 structure. These observations suggest that D10 and D2 co-aggregation could feature mixed amyloid fibrils, as previously observed in other systems^22^. We solved the structure of fibrils formed by purified recombinant D2-NT (**Figs. S5 and S6**). D2-NT fibrils differ from the D10-NT fibrils, but certain elements are conserved. The first seven residues of strand-1 adopt a very similar conformation to that observed in the D10-NT structure, as do the last five residues of strand-3 and the two strands pack together in a similar manner in both structures, with residue F73(D10)/F84(D2) packing against the equivalent Q47(D10)/Q58(D2) residue (**Fig. S7**).

However, in the D2-NT structure, strand-1 of PF-1 packs against strand-3 of PF-2, because the beta-helix that connects the two strands in the D10-NT structure reverses on itself, allowing the C-terminal region of D2 PF-1 to occupy location of the C-terminal region of PF-2 in the D10-NT structure. Notably, the sequence alterations between D10 and D2 behave as predicted above, with D2 residues L76, H78 and I80 remaining at positions facing the outside of the fibril structure, and D2 residues T75 and G83 adopting similar poses to V64 and A72 in the D10-NT structure (**Fig. S6E**). Finally, the disease-linked D2 mutant p.T61I occurs at a position facing the interior of the D2 PF fold (**Fig. S6E**), suggesting that this mutation, too, would result in an altered fibril structure.

Here we show for the first time that the low-complexity domains of D10 and D2 form amyloid fibrils in vitro and in vivo. Structures of D10 and D2 fibrils show that their core comprises the most highly conserved region between the two proteins. At present, we do not have structures of ex-vivo D10 or D2 fibrils, but despite potential differences, specific features of ex-vivo fibrils are often captured by in vitro fibril structures^23,24^. Indeed, the structures we obtained provide compelling explanations for the variable phenotypes associated with different D10 mutations and for the reported co-aggregation of D10 and D2 in vivo, supporting their relevance to disease and providing a previously unappreciated link between D10/D2 amyloid fibril formation and neurodegeneration. Finally, our results also predict that increased levels of WT D10 or D2 could be associated with amyloid formation and disease.

## Acknowledgements

This work was funded in part by National Institutes of Health (NIH) grants RF1AG066493 (DE), R35NS122209 (GM), U01NS134684 (NAS), F31AG077836 (CDM) and F31HL154651 (NMS), by MDA grant 961871-02 (HK), and by Project ALS 2021-01 (HK). The WCM NMR core is supported by NIH shared instrumentation grants S10OD028556 and S10OD016320. Some of the work was performed at the WCM Cryo-EM Core Facility, at the NYU Langone Health’s Cryo-Electron Microscopy Laboratory (RRID: SCR_019202), which is partially supported by the Laura and Isaac Perlmutter Cancer Center Support Grant NIH/NCI P30CA016087, and at the Simons Electron Microscopy Center at the New York Structural Biology Center, with major support from the Simons Foundation (SF349247). We acknowledge assistance from Clay Bracken and Emily Grasso with NMR data collection and processing, from Carl Fluck for cryo-EM data collection, and from Biao Qiu, Carl Fluck, Greg Alushin and Sjors Scheres for cryo-EM data processing.

## Competing Interests

The authors declare no competing interests.

## Data and Resource Availability Statement

NMR resonance assignments for D10-NT and D2-NT will be available in the BMRB databank under accession numbers 52550 and 52551, respectively, upon publication. PDB files for the D10-NT, S59L D10-NT and D2-NT fibrils will be deposited in the Protein Data Bank and cryo-EM maps will be deposited in EMDataResource upon publication. All other data and reagents used in this study are available upon request from the corresponding authors.

## Notes

### Competing Interest Statement

The authors have declared no competing interest.

