## Supplementary Figures and Online Methods for "Amyloid fibril structures link CHCHD10 and CHCHD2 to neurodegeneration"

**Figure S1: A**) Schematic of human CHCHD10, indicating the putative mitochondrial targeting sequence (MTS, red) situated at the N-terminus of the disordered N-terminal domain (blue) and the folded CHCH domain (black). **B**) NMR $^{15}N$-$^{1}H$ HSQC spectra of D10-NT. **C**) Secondary carbon chemical shifts of D10-NT.

**A**

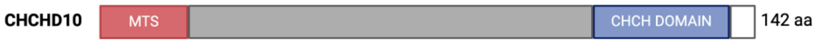


**B**

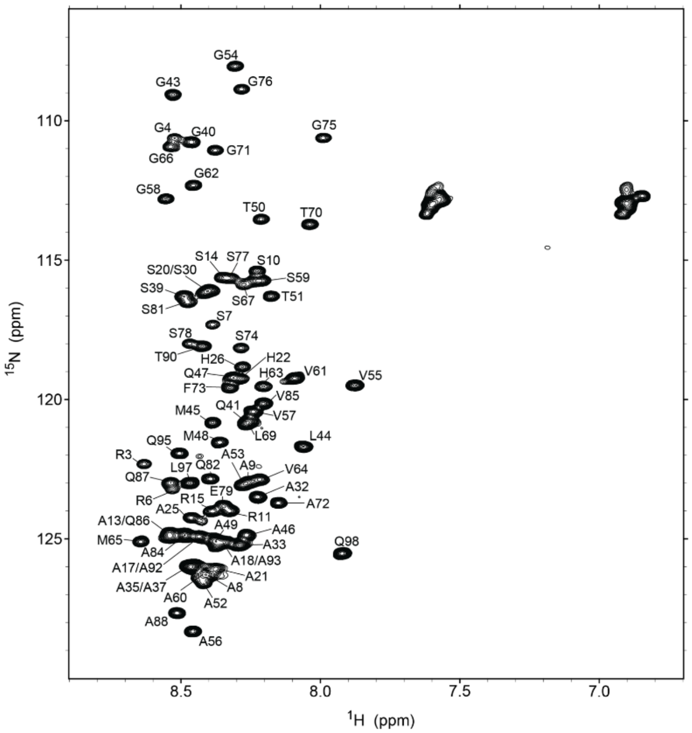


**C**

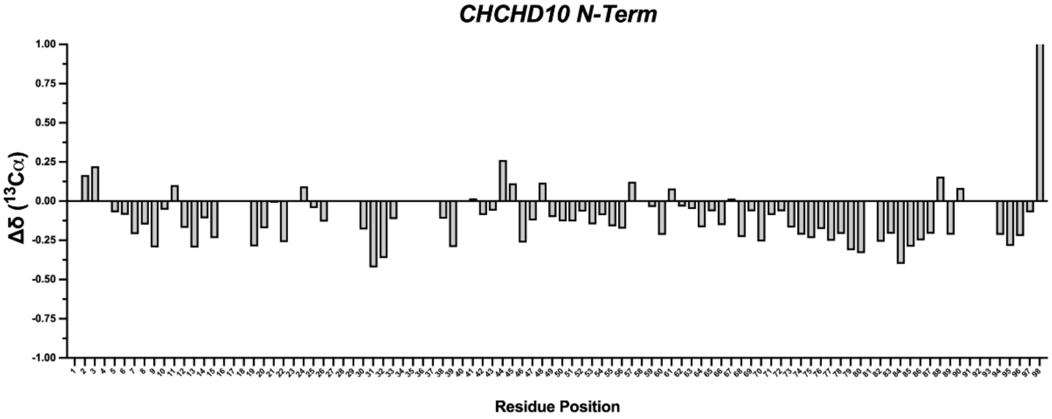

**Figure S2: A**) Sequence alignment of human and mouse D10 with residues S59 (human) and S55 (mouse) in red. **B**) Filter trap capture of extracted sarkosyl-insoluble aggregates from D10$^{WT}$, D10$^{S55L}$, and D10$^{KO}$ mouse hearts, immunoblotted (IB) for D10. **C**) TEM images of sarkosyl-insoluble fibrils extracted from D10$^{WT}$, D10$^{S55L}$, and D10$^{KO}$ mouse hearts immuno-gold labeled for D10. White arrows point to immuno-gold labelled D10. Red arrow heads point to amyloid-like fibrils on the exterior of a large clumped aggregate. Scale bar: 100 nm.

**A**

```
Identities:119/142(84%), Positives:126/142(88%), Gaps:4/142(2%)

Human CHCHD10  1    MPRGSRSAASRPASRPAAPSAHPPAHPPPSAAAPAPAPSGQPGLMAQMATTAAGVAVGSA  60
                    MPRGSRSAA+RP SRPA P AHPP    PSAAAPAPA  GQPGLMAQMA+TAAGVAVGSA
Mouse CHCHD10  1    MPRGSRSAAARPVSRPAPPPAHPP----PSAAAPAPAAPGQPGLMAQMASTAAGVAVGSA  56

Human CHCHD10  61   VGHVMGSALTGAFSGGSSEPSQPAVQQAPTPAAPQPLQMGPCAYEIRQFLDCSTTQSDLS  120
                    VGHVMGSALT AFSGG+SEP+QPAVQQAP   A  PLQMGPC+YEI+QFLDCSTTQSDL+
Mouse CHCHD10  57   VGHVMGSALTSAFSGGNSEPAQPAVQQAPARPASHPLQMGPCSYEIKQFLDCSTTQSDLT  116

Human CHCHD10  121  LCEGFSEALKQCKYYHGLSSLP  142
                    LCEGFSEALKQCKY HGLSSLP
Mouse CHCHD10  117  LCEGFSEALKQCKYNHGLSSLP  138
```

**B**

D10$^{WT}$ D10$^{S55L}$ D10$^{KO}$

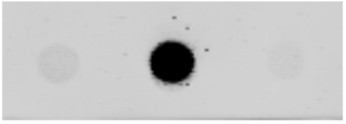

IB: CHCHD10

**C**

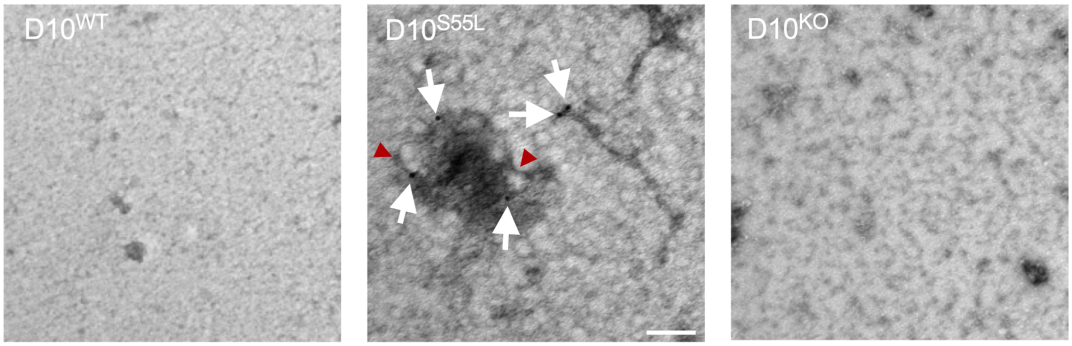

**Figure S3: A**) Representative cryo-EM micrograph of D10-NT fibrils. **B**) 2D class-averages (box size 256 pixels) of the major form of D10-NT fibrils. **C**) Map and model validations for D10-NT fibrils, including Fourier shell correlation (FSC) curves for the density map (blue), for the refined model versus full map (cyan), and for half maps for cross-validation (purple and pink). Black and dashed lines correspond to FSC values of 0.5 and 0.143, respectively.

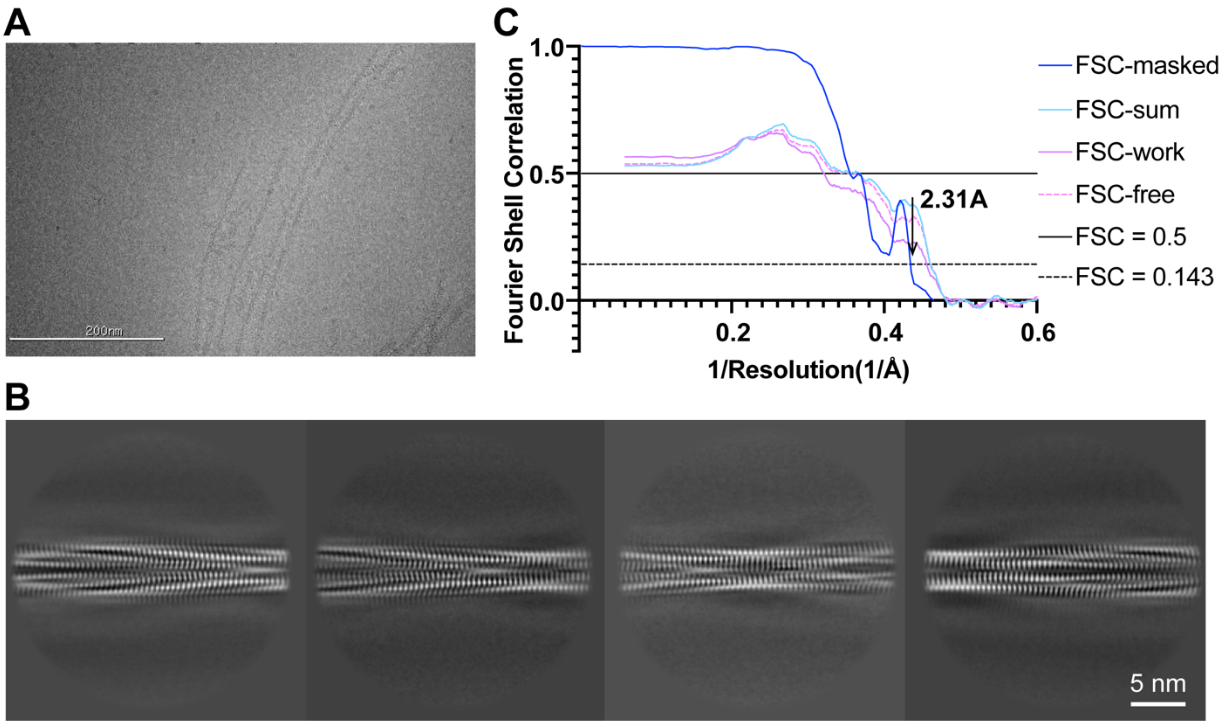

**Figure S4: A**) Representative cryo-EM micrograph of S59L D10-NT fibrils. **B**) 2D class-averages (box size 384 pixels) of S59L D10-NT short-crossover fibrils. **C**) Cryo-EM density map and atomic model of S59L D10-NT short fibrils. Three layers of the fibril structure are shown. **D**) Cryo-EM density map colored according to local resolution. **E**) Stick representation of a single layer of the S59L D10-NT ordered fibril core (PF1: G43-G75; PF2: L44-S74). Residue L59 is in red. **F**) Map and model validations for D10-NT fibrils, including FSC curves for the density map (blue), for the refined model versus full map (cyan), and for half maps for cross-validation (purple and pink). Black and dashed lines correspond to FSC values of 0.5 and 0.143, respectively.

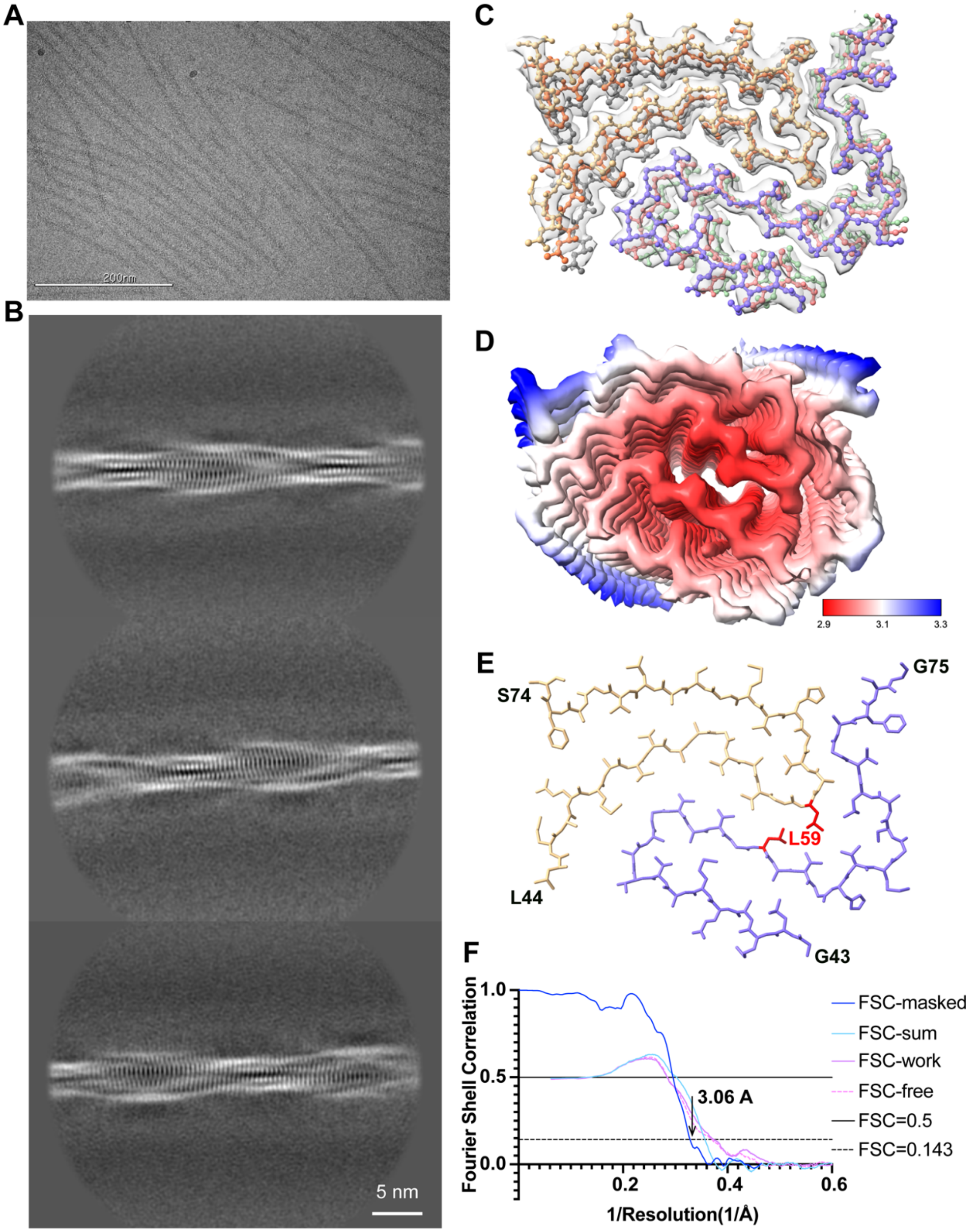

**Figure S5: A**) Schematic of human CHCHD2, indicating the putative mitochondrial targeting sequence (MTS, red) situated at the N-terminus of the disordered N-terminal domain (grey) and the folded CHCH domain (blue). **B**) Circular dichroism spectra of full-length D2 (black) and D2-NT (blue). **C**) Thioflavin T monitored aggregation of purified recombinant D2-NT. **D**) NMR $^{15}N$-$^{1}H$ HSQC spectra of D10-NT. **E**) Secondary carbon chemical shifts of D2-NT.

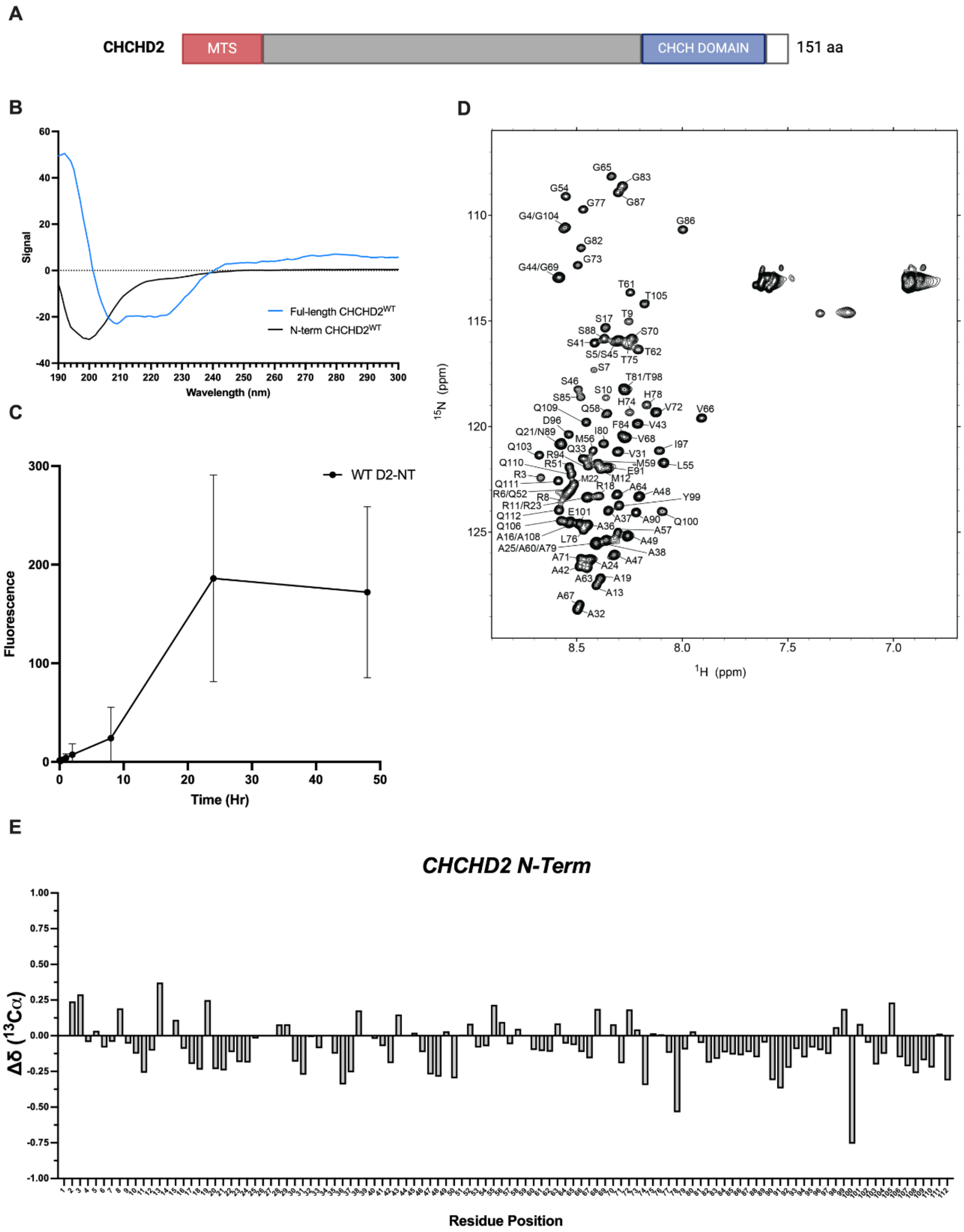

**Figure S6:** **A**) Representative cryo-EM micrograph of D2-NT fibrils. **B**) 2D class-averages (box size 640 pixels) of the major form of D2-NT fibrils. **C**) Cryo-EM density map and atomic model of D2-NT fibrils. Three layers of the fibril structure are shown. **D**) Cryo-EM density map colored according to local resolution. **E**) Stick representation of a single layer of the D2-NT ordered fibril core (G54-G86). Residues T75, L76, H78, I80 and G83, corresponding to D10 residues V64, M65, S67, L69 and A72, are in green. Residue T61 is in red. **F**) Map and model validations for D10-NT fibrils, including FSC curves for the density map (blue), for the refined model versus full map (cyan), and for half maps for cross-validation (purple and pink). Black and dashed lines correspond to FSC values of 0.5 and 0.143, respectively.

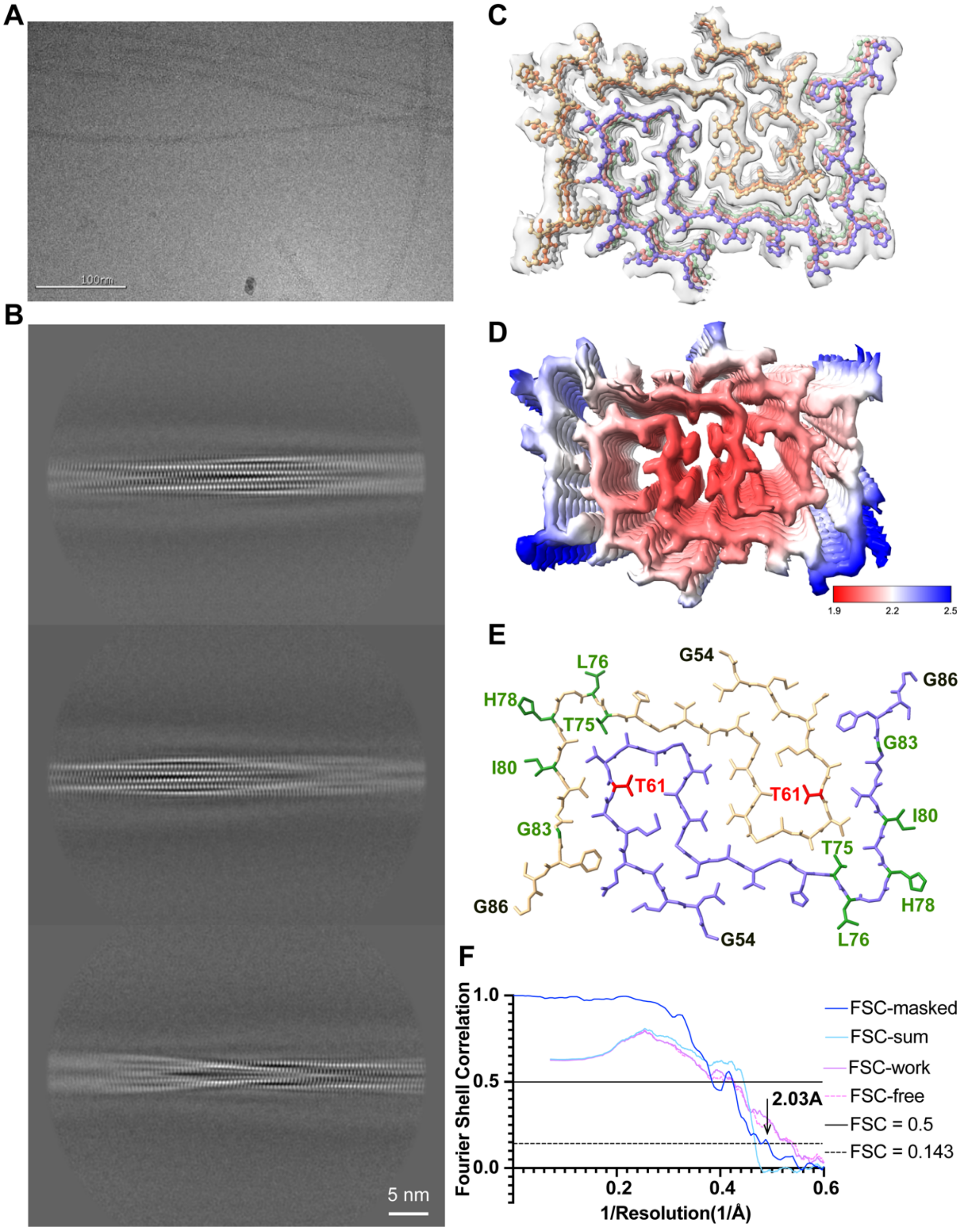

**Figure S7:** Secondary structure of WT D10-NT **(A)** and D2-NT **(B)** ordered fibril core structures showing the intra-molecular (D10-NT) vs. Inter-molecular (D2-NT) packing arrangement of strands 1 and 3 **(C,D)** Overlay of D10-NT and D2-NT structures, highlighting the similar conformations of the N-terminus of strand-1 (D10 residues G43-T51; D2 residues G54-T62) and the C-terminus of strand-3 (D10 residues A68-G75; D2 residues A79-G86) and of their detailed packing arrangement.

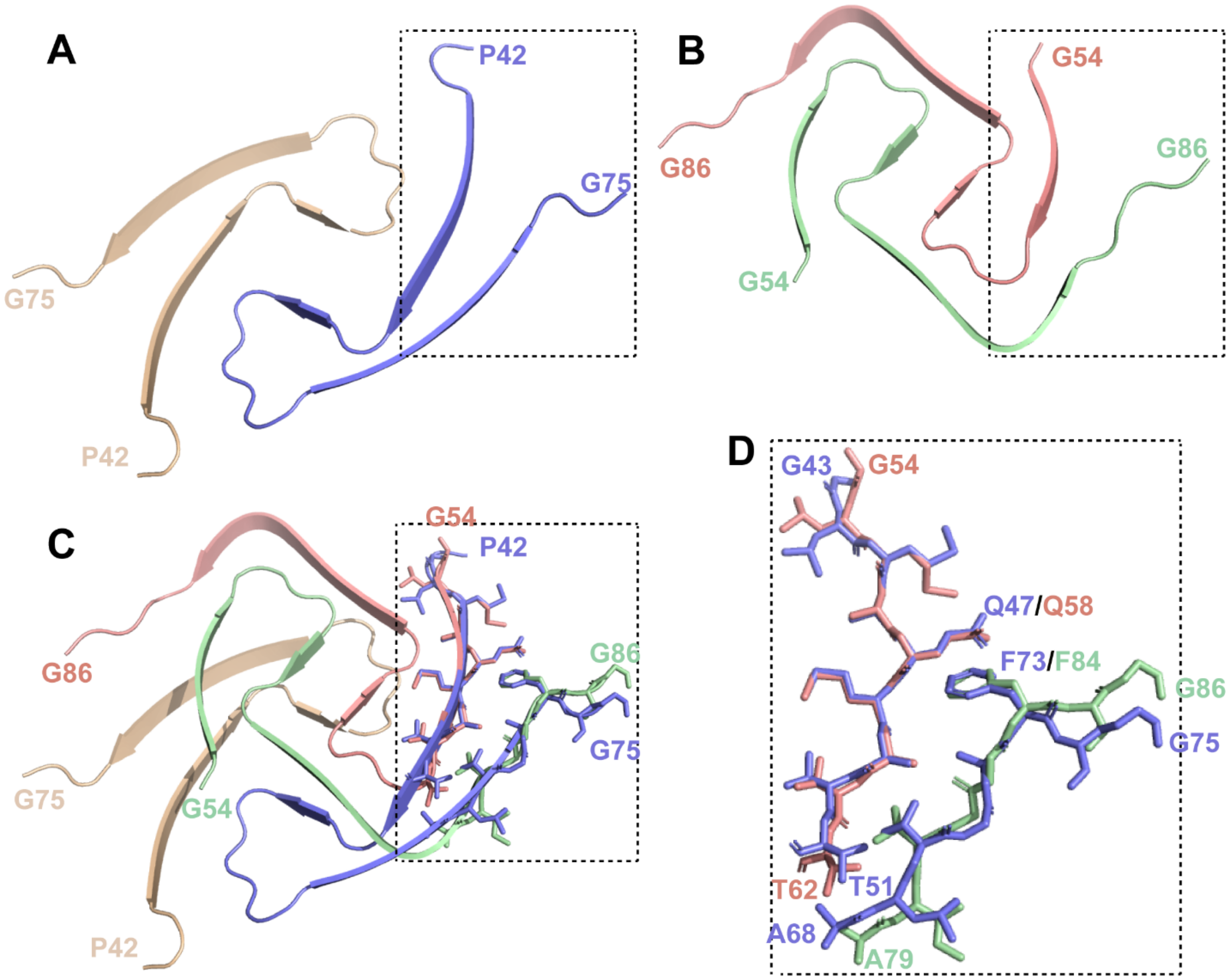

**Table 1. Statistics of cryo-EM data collection and refinement**

| | D10-NT | S59L D10-NT | D2-NT |
|---|---|---|---|
| PDB ID | | | |
| EMDB ID | | | |
| | **Data collection** | | |
| Magnification | 64,000 | 64,000 | 105,000 |
| Pixel size (Å) | 1.076 | 1.076 | 0.825 |
| Defocus Range (μm) | 0.9 to 2.2 | 0.8 to 2.5 | 0.6-2.5 |
| Voltage (kV) | 300 | 300 | 300 |
| Energy filter | 20 eV | 20 eV | 20 eV |
| Microscope/camera | Krios/K3 | Krios/K3 | Krios/K3 |
| Exposure time (s/frame) | 0.05 | 0.028 | 0.045 |
| Number of frames | 50 | 50 | 40 |
| Total dose ($e^{-}/Å^{2}$) | 51.82 | 59.653 | 48.4 |
| | **Reconstruction** | | |
| Micrographs | 8,278 | 4228 | 5892 |
| Manually picked fibrils | 5,167 | 1,543 | 5,435 |
| Box size (pixel) | 256 | 256 | 256 |
| Inter-box distance (Å) | 3.255 | 3.255 | 4.95 |
| Segments extracted (no.) | 1,902,596 | 658,614 | 1,496,493 |
| Segments after Class2D (no.) | 667,405 | 61,902 | 63,528 |
| Segments after Class3D (no.) | 602,449 | 19,650 | 23,351 |
| Resolution (Å) | 2.31 | 3.06 | 2.03 |
| Map sharpening B-factor ($Å^{2}$) | −56.6379 | -50.3765 | -23.67 |
| symmetry | pseudo-$2_1$ | C1 | C1 |
| Helical rise (Å) | 2.365 | 4.65 | 4.796 |
| Helical twist (°) | 178.845 | 3.88 | 1.34 |
| | **Atomic model** | | |
| Non-hydrogen atoms | 1,272 | 1,218 | 1,664 |
| Protein residues | 204 | 192 | 264 |
| Ligands | 0 | 0 | 0 |
| r.m.s.d. Bond lengths | 0.002 | 0.002 | 0.001 |
| r.m.s.d. Bond angles | 0.002 | 0.501 | 0.416 |
| MolProbity score | 0.825 | 1.42 | 1.48 |
| All-atom clashscore | 1.30 | 7.7 | 9.08 |
| Rotamer outliers (%) | 5.49 | 0 | 0 |
| Ramachandran Outliers (%) | 0 | 0 | 0 |
| Ramachandran Allowed (%) | 0 | 1.67 | 0 |
| Ramachandran Favored (%) | 100 | 98.33 | 100 |
| Ramachandran Disallowed (%) | 0 | 0 | 0 |

**Online Methods**

*Protein Expression and Purification:* Plasmids encoding human CHCHD10, CHCHD2, the CHCHD10 N-Terminus (residues 1-99) and the CHCHD2 N-Terminus (residues 1-114) preceded by an N-terminal 6x-His-SUMO tag were procured from Twist Biosciences. The p.S59L CHCHD10 mutant was generated using an In-Fusion Cloning kit (Takara Bio) and confirmed by DNA sequencing (Genewiz). Recombinant proteins were expressed in E. coli BL21/DE3 cells (Novagen) grown in either LB Broth or M9 minimal media containing 15N-labeled ammonium chloride (1 g/L) or 15N-labeled ammonium chloride and 13C-labeled D-glucose (2 g/L) at 37 °C (275 rpm) induced with 1 mM IPTG (Isopropyl b-D-1-thiocalactopyranoside) at OD 600nm of 0.6-0.8. 4 hours post induction, cells were harvested via centrifugation at ca. 10,500 g at 4 °C for 15 mins. Cell pellets (stored at -20 °C overnight) were resuspended in 50 mL lysis buffer (350mM NaCl, 20 mM Imidazole, 20 mM Tris pH 8.0, 1mM PMSF (phenylmethylsulfonyl fluoride), 1 mM EDTA and 3 mM bME) and lysed using an EmulsiFlex-C3 (AVESTIN, Ontario, Canada), followed by centrifugation at ca. 40,000 g for 1 hour to remove cellular debris. The supernatant was loaded onto a Ni-NTA column equilibrated using 350 mM NaCl, 20 mM Imidazole, 20 mM Tris pH 8.0, 3mM bME, washed with the same buffer and the SUMO-tagged protein was eluted using 350 mM NaCl, 250 mM Imidazole, 20 mM Tris pH 8.0, 3 mM 2-mercaptoethanol. Protein-containing fractions were pooled and cleaved overnight using SUMO protease (added to final concentration ca. 1 µM), followed by dialysis against 150 mM NaCl, 20 mM Tris pH 8.0, and 1mM DTT and loaded again onto a Ni-NTA column. The cleaved N-terminal constructs were collected in the flowthrough and further purified over a 5 mL HiTrapTM SP HP column on an AKTA Pure Protein Purification System (GE). Purified constructs were then exchanged into diH2O using a PD-10 Column (Cytiva, Marlborough, MA) for lyophilization.

*Circular Dichroism (CD) Spectroscopy:* CD measurements were performed on an AVIV 410 CD spectropolarimeter. Spectra were obtained from 300-190 nm at 25 °C after a two-minute temperature equilibration with a wavelength step of 1 nm, an averaging time of 5 seconds, 1 scan per sample and a cell pathlength of 0.02 cm (Starna, Atascadero, CA). Backgrounds were collected and subtracted from all spectra. Final construct concentrations ranged from ca. 50-100 µM as assessed by 1D proton NMR with DSS as an internal standard. The dearth of aromatic residues in all of the polypeptides used made reliable determination of absolute protein concentrations exceedingly difficult. Therefore, CD data are presented in millidegrees and were not converted to mean residue molar ellipticity.

*Thioflavin T Aggregation Reaction:* Monitoring in vitro CHCHD10 and CHCHD2 aggregation using Thioflavin T was adopted from a report on tau aggregation[1]. Lyophilized protein was brought up to stock concentrations of ~16 µM in aggregation assay buffer (20 mM Tris pH 7.4, 100 mM NaCl, 1 mM EDTA and 1 mM DTT) and filtered using centrifugation and a 100 kDa (AMICON) at room temperature for 10 mins at ca. 15,000 x g before initializing the reaction to remove any initial aggregate formation. A 3 mM Thioflavin T stock was prepared in aggregation assay buffer and filtered using 0.2 µm filter (Pall). Aggregation was induced using an eppendorf Thermomixer R at 37 °C shaking at 1000 rpm. Data was collected in triplicates at designated time intervals using a microplate fluorescence reader (Molecular Devices). The excitation and emission wavelengths were 450 nm and 510 nm respectively.

*Extraction of sarcosyl-insoluble fractions:* Sarcosyl-insoluble fractions were obtained as previously described[2] with minor adjustments. 20 mg of heart tissue was homogenized in 1 ml of extraction buffer (10 mM Tris-HCl, pH 7.4, 0.8 M NaCl, 10% sucrose, 1 mM EGTA, 2% sarcosyl,1X cOmplete protease inhibitor) using a Tissue-Tearor (BioSpec) for 1 min on ice. Homogenized tissue was incubated at 37 °C for 1 hour, then centrifuged at 10,000 x g for 10 min at 4 °C to remove debris. Supernatant was collected and spun at 100,000 x g for 1 hour at 4 °C. Pellet was resuspended in 150 µl extraction buffer and centrifuged at 3,000 x g for 5 min at 4 °C. Supernatant was raised to 1 ml in 50 mM Tris-HCl, pH 7.4, 150 mM NaCl, 10% sucrose, 0.2% sarcosyl, and 1X cOmplete protease inhibitor, then centrifuged at 100,000 x g for 30 min at 4 °C. Sarcosyl-insoluble pellet was resuspended in 50 µl 20 mM Tris-HCl, pH 7.4, 50 mM NaCl and was used for filter trap assay, immunogold-labelling and electron microscopy. Of note, N-Lauroylsarcosine (sarcosyl) sodium salt (Sigma) was used for these experiments.

*Filter Trap Assay:* Insoluble protein aggregates were detected by filter trap assay as previously described[3]. Briefly, 1 µl of the final sarcosyl-insoluble fraction was loaded onto a Bio-Dot Microfiltration apparatus (Bio-Rad) containing a cellulose acetate membrane (0.2 µm pore diameter, Whatman). Vacuum was applied to pass samples through the membrane, which was then was washed with 1% Tween-20 in PBS. Trapped proteins were detected with rabbit anti-CHCHD10 (Abcam Ab121196). Blots were then imaged using Clarity Western ECL Blotting Substrates (Bio-Rad) and imaged on ChemiDoc Touch (Bio-Rad).

*Immunogold preparation of sarkosyl-insoluble fractions for EM*: 5 µl of sarkosyl-insoluble sample was applied to a formvar-carbon coated grid and allowed to settle for 2 min. Grids were inverted onto a 100 µl drop of Aurion Blocking buffer for the secondary antibodies (Aurion, Electron Microscopy Sciences) for 15 min. Grids were incubated on 100 µl drops of primary rabbit-anti-CHCHD10 (Abcam Ab121196) in PBS-c (PBS + 0.1% BSA-c, Aurion) for 1 hour in a human box at room temperature. Grids were washed 5 times in PBS-c for 5 min. Grids were incubated on 100 µl drops of gold-tagged secondary antibody (Aurion) for 1 hour in a humid box, then washed 3 times for 3 min in PBS-c and two times for 3 minutes in diH2O. Grids were fixed with 2.5% buffered glutaraldehyde (Sigma) and washed 3 times for 1 min in diH2O. Negative stain was applied with 1.5% (aq) uranyl acetate (Sigma), blotted and dried. EM imaging used a JEOL JEM 1400 transmission electron microscope operated at 100 kV and equipped with an Olympus-SIS Veleta side-mount 2K × 2K digital camera.

*Fibroblast imaging:* Fibroblasts from individuals with CHCHD10 p.R15L mutation (Columbia University, NY) and cultured on glass coverslips (Electron Microscopy Sciences 50-949-008) in DMEM with 10 % FBS to 80% confluency before fixation with 4% paraformaldehyde. Fixed fibroblasts were permeabilized with 0.1% Triton X-100 in PBS, blocked, and then stained with anti-CHCHD10 primary antibody (ProteinTech 25671-1-AP, Rabbit polyclonal, 1:500 dilution) and Thioflavin S to detect amyloid deposits. Thioflavin S (Sigma-Aldrich T1892) was prepared by dissolving it in 50% ethanol to 0.05% w/v and filtering the solution through a 0.2 µM PES filter (Thermo Scientific 725-2520). For Thioflavin S staining, fibroblasts were incubated with this solution for 5 minutes at room temperature, followed by sequential washes in 50%, 80% and 95% ethanol then washed several more times in Milli-Q $H_2O$. Fibroblasts were then imaged by laser scanning confocal microscopy (85 µM pinhole) using a Leica SP5 system equipped with a 40x oil objective. The Thioflavin S staining was captured using 488 nm excitation and 500-538 nm PMT detection. CHCHD10 and Thioflavin S overlays were created using ImageJ2 (FIJI distribution).

*Negative Stain Electron Microscopy:* 5 µl of aggregated D10-NT sample was applied to a formvar-carbon coated grid and allowed to settle for 1 min. Grids were washed with diH2O 3 times, stained with 1.5% (aq) uranyl acetate (Sigma), incubated with stain for 1min, washed with diH2O, blotted with filter paper and dried. EM imaging used a JEOL JEM 1400 transmission electron microscope operated at 100 kV and equipped with an Olympus-SIS Veleta side-mount 2K × 2K digital camera.

*Solution state Nuclear Magnetic Resonance (NMR) Spectroscopy:* CHCHD10 and CHCHD2 constructs were prepared in NMR Buffer (100 mM NaCl, 10 mM Na2HPO4, pH 6.8) at concentrations ranging from ca. 50-150 µM in 5 mm NMR tubes. Relative protein concentrations were corroborated by 1D proton NMR using 4,4-dimethyl-4-silapentane-1-sulfonic acid (DSS) as an internal standard. $^{1}H$-$^{15}N$ HSQC spectra were collected on a Bruker AVANCE 500-MHz spectrometer equipped with a Bruker TCI cryoprobe at 10 °C with 1024 complex points in the $^{1}H$ dimension and 214 complex points in the $^{15}N$ dimension using spectral widths of 18 PPM ($^{1}H$) and 30 PPM ($^{15}N$). NMR spectra were processed using NMRpipe and analyzed using NMRFAM-sparky 3.115 and NMRbox. Backbone resonance assignments were made using standard triple resonance experiments collected on a Bruker Avance III spectrometer at 800 MHz.

*Cryo-EM data collection:* Fibrilization of D10-NT, S59L D10-NT or D2-NT was performed as described above by incubating proteins at 37 °C, shaking at 1200 rpm, and monitoring reaction aliquots for Thioflavin T fluorescence. 3.5 µL aliquot of D10-NT fibrils aggregated for 5 days at an initial monomer concentration of ca. 80 µM, S59L D10-NT fibrils aggregated for 22 hours at an initial monomer concentration of ca. 40 µM, or D2-NT fibrils aggregated for 7 days at an initial monomer concentration of ca. 40 µM, was applied to a glow-discharged holey copper grid (Quantifoil R1.2/1.3, 300 mesh) and incubated for 30 s. The grids were blotted for 3 s under 100% humidity at 20 °C and plunge-frozen into liquid ethane using a Vitrobot Mark IV (FEI, Thermo). The data sets for D10-NT fibrils and S59L D10-NT fibrils were collected using Leginon[4] at the New York Structural Biology Center. The data set for D2-NT fibrils was collected using Leginon[4] at New York University Langone Health's Cryo-EM Laboratory. The detailed parameters of cryo-EM data collection are provided in Table 1.

*Cryo-EM image processing and helical reconstruction:* All datasets were processed using Relion 4.0.0 [5,6]. Frame stacks were aligned with dose-weighting using MotionCorr2 [7], and the contrast transfer function (CTF) was estimated using CTFFIND4.1 [8]. Helical reconstruction was performed in RELION 4.0.0 [5,6,9]. Fibrils were manually picked and segments were extracted using a large box size, typically 768 pixels. Reference-free 2D classification was conducted for several rounds to obtain homogeneous subsets. Helical pitch was estimated from the 2D classes using ImageJ. Further re-extraction to a smaller box size, typically 384 pixels, and 2D classification was conducted and the resulting 2D classes were used to generate initial models using the relion_helix_inimodel2d program[5]. Segments were re-extracted to smaller box size (256 pixels) and subjected to 3D auto-refinement without a mask. A mask was then created and a 3D auto-refinement was conducted with mask. A further 3D auto-refinement was conducted with optimization of helical twist and rise. C1 with pseudo-$2_1$ screw symmetry and C2 symmetry were also separately imposed in the refinements to test if there was further improvement of map. The refined unfiltered half-map was postprocessed using a soft mask spanning 20% of the box along the helical axis. The particles were then Bayesian polished and subjected to a further round of 3D auto-refinement to improve the resolution. 3D classification and another round of 3D auto-refinement was performed to the best class. The final reconstruction was sharpened by applying the standard post-processing procedure. All reconstruction details are shown in Table 1.

*Cryo-EM model building and refinement*: Atomic models were built de novo and modified by COOT[10]. Models were iteratively refined using the Real-space refinement package in PHENIX[11] and validated using MolProbity[12]. For cross validation, all atoms in the refined models were randomly displaced by an average of 0.3 Å, and each resulting model was refined against the first unfiltered half-map obtained from processing (FSC-work). FSC curves were then calculated between the FSC-work model and the second unfiltered half-map obtained from processing

(FSC-free), and between the refined models and the full-map (FSC-sum). The structural figures were prepared in UCSF ChimeraX[13] and PyMOL (DeLano Scientific). The detailed parameters of atomic model building are provided in Table 1.
